# Chromosome-level genome assembly and annotation of the crested gecko, *Correlophus ciliatus*, a lizard incapable of tail regeneration

**DOI:** 10.1101/2024.09.28.615630

**Authors:** Marc A Gumangan, Zheyu Pan, Thomas P Lozito

**Affiliations:** Department of Orthopaedic Surgery, Keck School of Medicine, University of Southern California, 1540 Alcazar St, Los Angeles, CA 90089, USA; Department of Stem Cell Biology and Regenerative Medicine, Keck School of Medicine, University of Southern California, 1425 San Pablo St, Los Angeles, CA 90033, USA

**Keywords:** Gekkota, crested gecko, Correlophus ciliatus, genome sequencing, assembly

## Abstract

The vast majority of gecko species are capable of tail regeneration, but singular geckos of *Correlophus, Uroplatus*, and *Nephrurus* genera are unable to regrow lost tails. Of these non-regenerative geckos, the crested gecko (*Correlophus ciliatus*) is distinguished by ready availability, ease of care, high productivity, and hybridization potential. These features make *C. ciliatus* particularly suited as a model for studying the genetic, molecular, and cellular mechanisms underlying loss of tail regeneration capabilities. We report a contiguous genome of *C. ciliatus* with a total size of 1.65 Gb, a total of 152 scaffolds, L50 of 6, and N50 of 109 Mb. Repetitive content consists of 40.41% of the genome, and a total of 30,780 genes were annotated. Assembly of the crested gecko genome provides a valuable resource for future comparative genomic studies between non-regenerative and regenerative geckos and other squamate reptiles.

**Findings:** We report genome sequencing, assembly, and annotation for the crested gecko, *Correlophus ciliatus*.

## Introduction

The crested gecko, *Correlophus ciliatus*, is a lizard species endemic to New Caledonia distinguished by eye and head projections/spines (Fig. 1 A, B) and the inability to regenerate amputated tails (Fig. 1 C, D). The chromosomes of *C. ciliatus* are typical of most New Caledonian geckos and exhibit a biarmed, acrocentric 2n=38 karyomorph (Fig. 1 E) [1, 2]. Crested geckos readily adapt to captivity; as a nocturnal, omnivorous species from a mild, tropical climate, *C. ciliatus* thrives at “room temperatures” and does not require the expensive lighting or insect diets obligatory in the maintenance of many other lizard species. Since *C. ciliatus* is able to breed nearly year-round without seasonal simulations, this species is also one of the most straight-forward and productive to breed in captivity.

**Figure 1.**
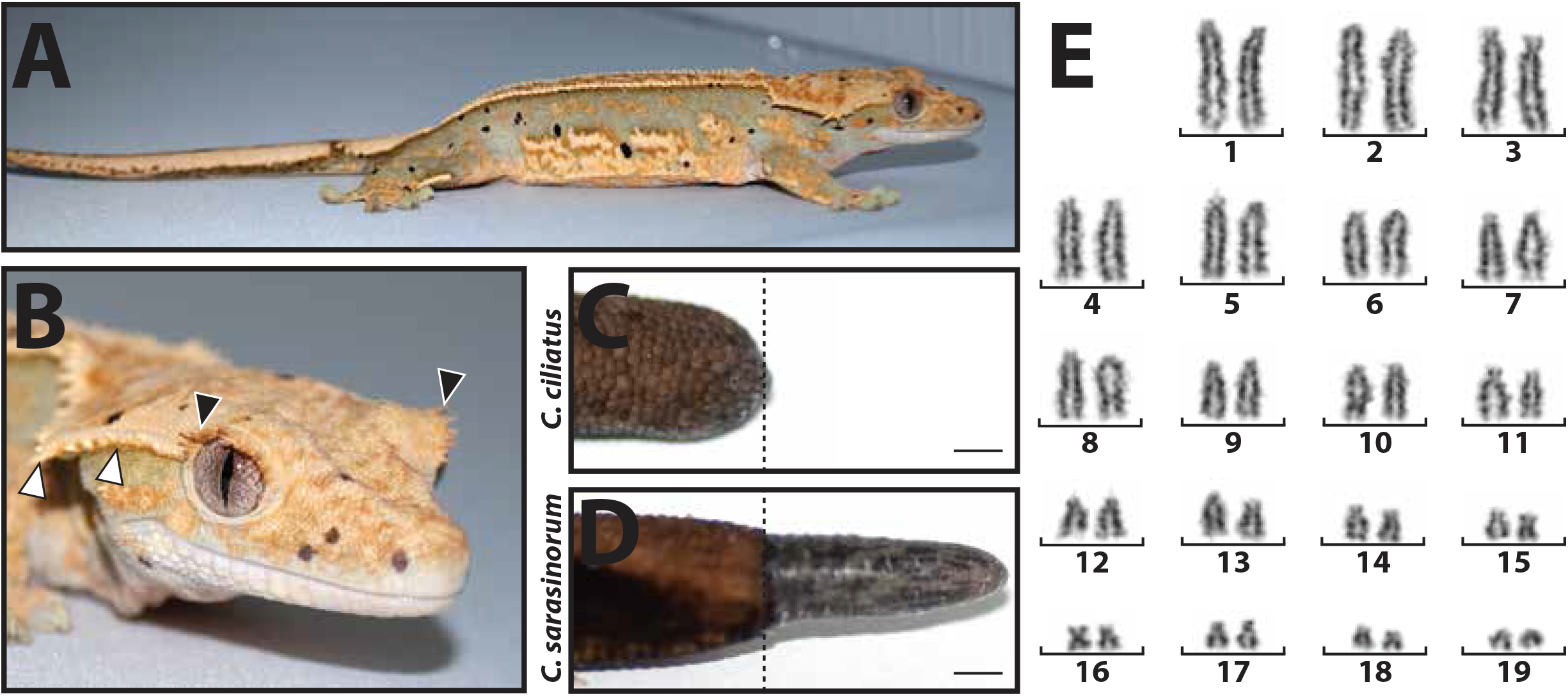
**(A)** Example of a crested gecko (*Correlophus ciliatus*). **(B)** *C. ciliatus* head detail showing head crest spines (white arrow heads) and “eyelash” spines (black arrow heads). **(C, D)** Representative gross anatomy images of *C. ciliatus* and *C. sarasinorum* tails 26 days post-amputation. *C. ciliatus* tails do not regenerate like those of related gecko species, including *C. sarasinorum*. Bar = 0.25 cm. **(E)** *C. ciliatus* karyotype (2n=38). Karyotype was prepared from *C. ciliatus* embryonic fibroblasts by the Molecular Cytogenetics Laboratory, Department of Veterinary Integrated Biosciences, Texas A&M University.

*C. ciliatus* is one of only fourteen described gecko species (over 1,850 total) that has lost the ability to regenerate amputated tails (Supplementary Material Table 1). Of these non-regenerative gecko species, only *C. ciliatus* is readily available within the American and European pet hobbies. Furthermore, *C. ciliatus* is the only non-regenerative lizard species capable of hybridizing with regenerative relatives, specifically *C. sarasinorum, Mniarogekko chahoua*, and *Rhacodactylus auriculatus*. Currently, all other gecko species with sequenced genomes are capable of tail regeneration (personal observation by T. P. L.) [3-7]. The goal of studying a non-regenerative gecko towards identifying gene regions involved in tail regrowth is a main driver for sequencing the *C. ciliatus* genome. With its ease of care, high productivity, and options for hybridizations, the crested gecko is the ideal model lizard for studying the genetic mechanisms involved in loss of tail regeneration capabilities.

**Table 1.** General statistics of the raw sequencing reads used for C. ciliatus assembly.

| Sample | Library Type | Sequencing Type | Platform | Number of Reads | Coverage |
| --- | --- | --- | --- | --- | --- |
| PacBio CCS | Long Reads | WGS | PacBio Sequel II | 13,182,509 | 62x |

## Methods

### Sample Collection, PacBio Sequencing and Assembly

Gecko housing, handling, and sample collections were performed according to the guidelines of the Institutional Animal Care and Use Committee at the University of Southern California (protocol 20992). Genomic DNA was obtained from a single whole female *C. ciliatus* embryo collected from a two-month-old egg incubated at 23°C. The Qiagen Midi Prep Kit was used for the DNA extraction from 94mg of ground embryo, and approximately 100 ug of high molecular weight (HMW) DNA was obtained. Genomic DNA was sequenced using the PacBio Sequel II platform (Table 1). 185.8 gigabase-pairs of PacBio CCS reads were used as inputs to Hifiasm v0.15.4-r347 [8] with default parameters. To estimate the genome size of *C. ciliatus*, k-mer analysis was conducted on the PacBio CCS read using a range of k values (17, 19, 21, 23, 25, 27, 29 and 31). The estimated genome size was calculated by: (total number of kmers – erroneous kmers) divided by homozygous peak depth, following the methods of Cai *et al* [9]. (Note: Minimum coverage was defined as the depth of the first trough in the k-mer frequency distribution. K-mers that fell under this minimum coverage were considered erroneous.) Jellyfish v2.2.10 [10] was used to calculate the k-mer frequency using the -C parameter, and GenomeScope v1.0.0 [11] was then used to estimate heterozygosity.

BLAST v2.9.0 [12] results of the Hifiasm output assembly against the NCBI Nucleotide Database were used as inputs for blobtools v1.1.1 [13]. Scaffolds identified as possible contamination (sequences within genomic data that originate from sources other than the intended target organism, *C. ciliatus*) were removed from the assembly. Blobtools revealed contamination from Actinobacteria (2 contigs, 15 Mb) (Fig. 2). Finally, purge_dups v1.2.5 [14] was used to remove haplotigs and contig overlaps.

**Figure 2.**
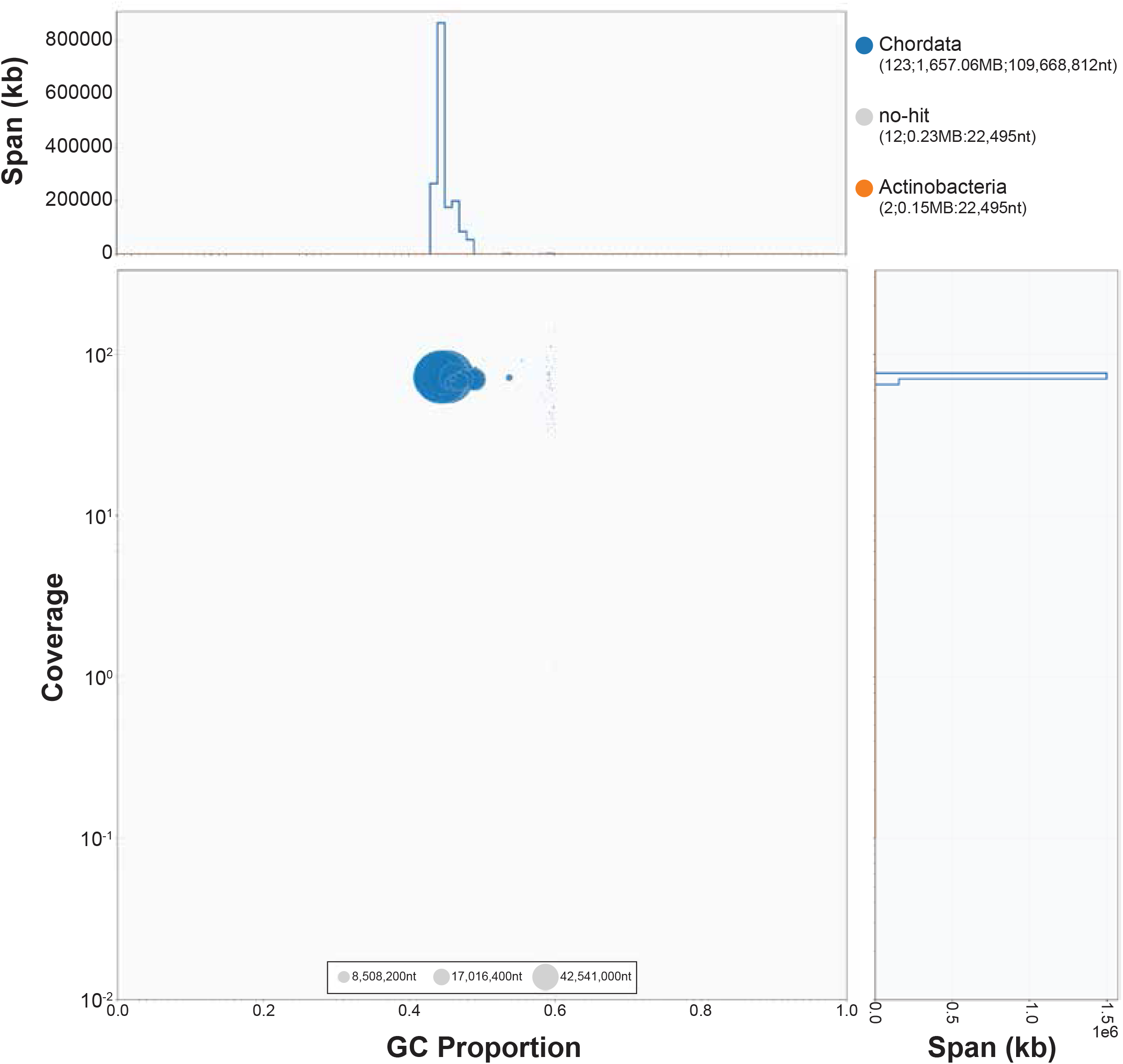
Blobtools plot showing taxonomic categories for different scaffolds (blue: Chordata, grey: ‘no hits’, orange: Actinobacteria), scaffold wide coverage, and GC content. Scaffolds under Actinobacteria were removed from the final genome assembly.

Dovetail Omni-Libraies were constructed to scaffold initial Hifiasm assemblies. For each Dovetail Omni-C library, nuclear chromatin was fixed with formaldehyde and extracted. Following digestion with DNAse I, chromatin ends were repaired and ligated to biotinylated bridge adapters followed by proximity ligation of adapter-containing ends. After proximity ligation, crosslinks were reversed, and the DNA was purified. Purified DNA was treated to remove free biotin that was not incorporated into ligated DNA fragments. Sequencing libraries were generated using NEBNext Ultra enzymes and Illumina-compatible adapters. Biotin-containing fragments were isolated using streptavidin beads before PCR enrichment of each library. Libraries were sequenced on an Illumina HiSeqX platform to produce approximately 30x sequence coverage.

The draft de novo assembly produced by Hifiasm and Dovetail OmniC library reads were input into HiRise [15], a software pipeline designed specifically for using proximity ligation data to scaffold genome assemblies. Dovetail OmniC library sequences were aligned to the draft assembly using bwa v0.7.17 [16]. The separations (genomic distances between pairs of reads that map within draft scaffolds) of Dovetail OmniC read pairs that mapped within draft scaffolds were analyzed by HiRise to produce a likelihood model for genomic distance between read pairs. This model was used to identify and break putative misjoins (erroneous link between two contigs), to score prospective joins, and make joins above a threshold.

### Repeat Content

De novo-based methods were used to identify transposable elements and other repetitive elements. Repetitive content within the *C. ciliatus* genome was predicted with RepeatModeler v2.0.1 [17]. Repetitive elements were masked using RepeatMasker v4.1.0 [18].

### Gene Annotation and BUSCO Analysis

Coding sequences from *Anolis carolinensis, Gekko japonicus, Pogona vitticeps, Salvator merianae*, and *Zootoca vivipara* were used to train the initial ab initio model for *C. ciliatus* using the AUGUSTUS software v 2.5.5 [19]. Six rounds of prediction optimization were performed with AUGUSTUS. The same coding sequences were also used to train a separate ab initio model for *C. ciliatus* using SNAP v2006-07-28 [20]. Total RNA was extracted from a single whole, two-month-old, female *C. ciliatus* embryo using the QIAGEN RNeasy Plus Kit following manufacturer protocols. Total RNA was quantified using Qubit RNA Assay and TapeStation 4200. Prior to library prep, DNase treatment was performed, followed by AMPure bead clean up and QIAGEN FastSelect HMR rRNA depletion. Library preparation was done with the NEBNext Ultra II RNA Library Prep Kit following manufacturer protocols. Then these libraries were run on the NovaSeq6000 platform in 2 x 150 bp configuration (Table 2). RNA-Seq reads were mapped onto the genome using the STAR alignment software v2.7 [21], and intron hints were generated with the bam2hints tools within the AUGUSTUS software. MAKER v3.01.03 [22], SNAP and AUGUSTUS (with intron-exon boundary hints provided from bam2hints) were then used to predict gene identities in the repeat-masked reference genome. To help guide the prediction process, Swiss-Prot peptide sequences from the UniProt database [23] were downloaded and used in conjunction with the protein sequences from *A. carolinensis, G. japonicus, P. vitticeps, S. merianae, Z. vivipara* to generate peptide evidence in the MAKER pipeline. Only gene identities that were predicted by both SNAP and AUGUSTUS softwares were retained in final gene sets.

**Table 2.**
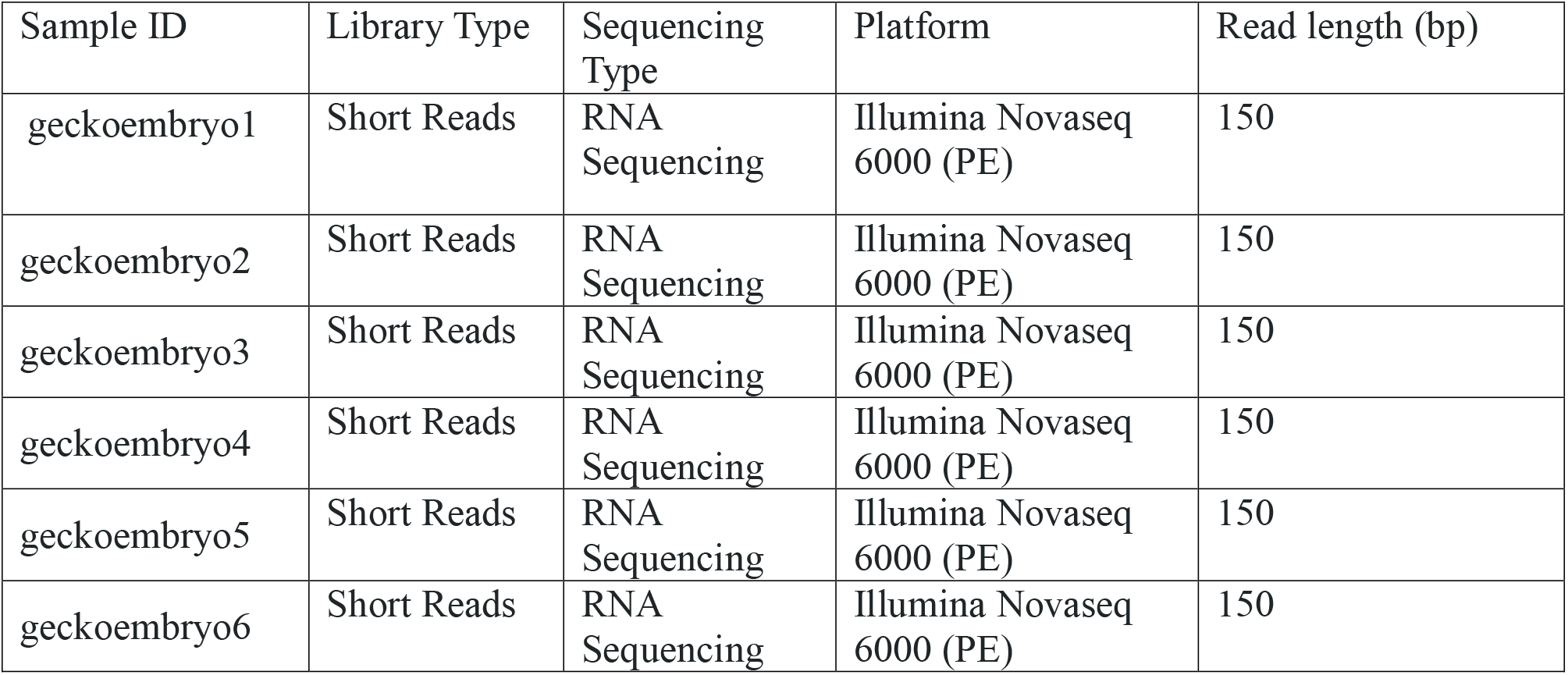
Summary of RNA-Seq samples statistics.

## Results and Discussion

The total assembly size is 1.65 Gb, with a GC content of 45% (Table 3). The estimated genome size by k-mer analysis was 1.52 Gb (Table 3). It is worth noting the approximately two-fold difference between the coverage of the heterozygous peak at 50X, and the homozygous peak at 100X (Fig. 3). This indicates a high level of heterozygosity in the genome. The rate of heterozygosity estimated by GenomeScope was approximately 0.51% (Table 4). The contig/scaffold N50 is 109 Mb, and the largest scaffold had a length 169 Mbp (Table 3). 99.54% (1653 Mbp) of the total assembly was scaffolded into 19 chromosome length scaffolds (Fig. 4). This number of chromosomal scaffolds is consistent with the number of haploid chromosomes observed in the *C. ciliatus* karyotype (Fig. 1 E).

**Table 3.** *C. ciliatus* genome assembly statistics.

|  | Contig | Scaffold |
| --- | --- | --- |
| Largest scaffold length (bp) | 169,053,634 | 169,053,634 |
| N90 (bp) | 26,447,670 | 51,189,016 |
| N50 (bp) | 109,210,969 | 109,210,969 |
| L90 | 6 | 19 |
| L50 | 6 | 14 |
| Number > 1kbp | 163 | 152 |
| Number of N's per 100 kbp | 0.00 | 0.07 |
| GC Content (%) | 45 | 45 |
| Total Size (bp) | 1,653,058,530 | 1,653,059,630 |

**Table 4.** Estimation of genome size and heterozygosity of *C. ciliatus* assembly.

| k | Total number of k-mers | Minimum coverage (X) | Number of erroneous k-mers | Homozygous peak | Heterozygous peak | Estimated genome size (Gb) | Estimated heterozygosity (%) |
| --- | --- | --- | --- | --- | --- | --- | --- |
| 17 | 160830997409 | 23 | 1309311311 | 110 | 55 | 1.45 | 0.46 |
| 19 | 163207828191 | 21 | 2876102102 | 108 | 53 | 1.48 | 0.51 |
| 21 | 164978566184 | 21 | 3624290072 | 107 | 53 | 1.51 | 0.52 |
| 23 | 166415784027 | 21 | 4147151401 | 107 | 53 | 1.52 | 0.52 |
| 25 | 167625136836 | 21 | 4629779608 | 107 | 52 | 1.52 | 0.52 |
| 27 | 168655291501 | 21 | 5100636735 | 106 | 52 | 1.54 | 0.51 |
| 29 | 169546294955 | 21 | 5566480734 | 106 | 52 | 1.55 | 0.50 |
| 31 | 170331212990 | 21 | 6029874685 | 106 | 52 | 1.55 | 0.50 |

**Figure 3.**
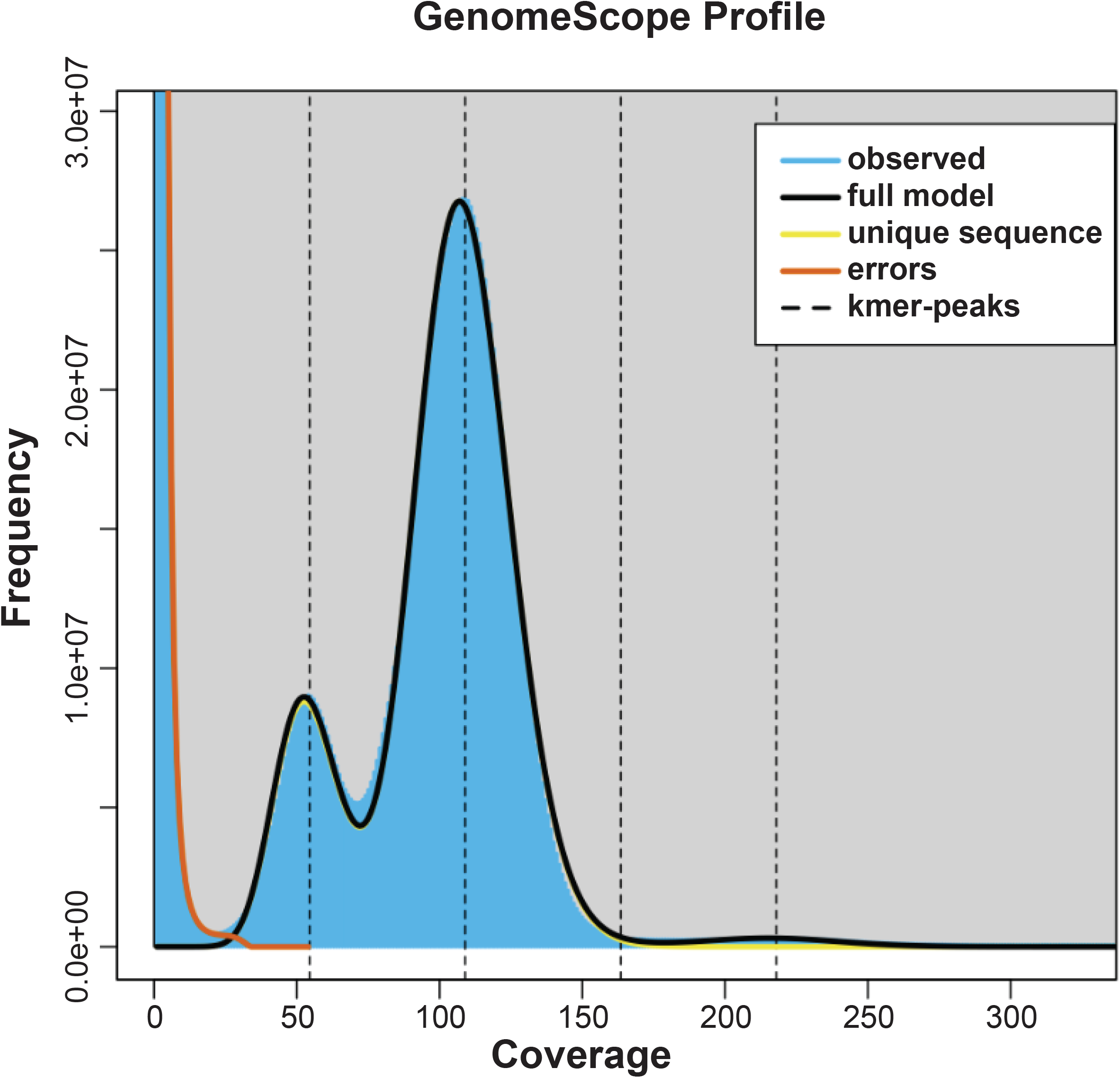
21-mer frequency distribution of the *C. ciliatus genome*. The first peak at 53X is the heterozygous peak, while the second peak at 107X is the homozygous peak.

**Figure 4.**
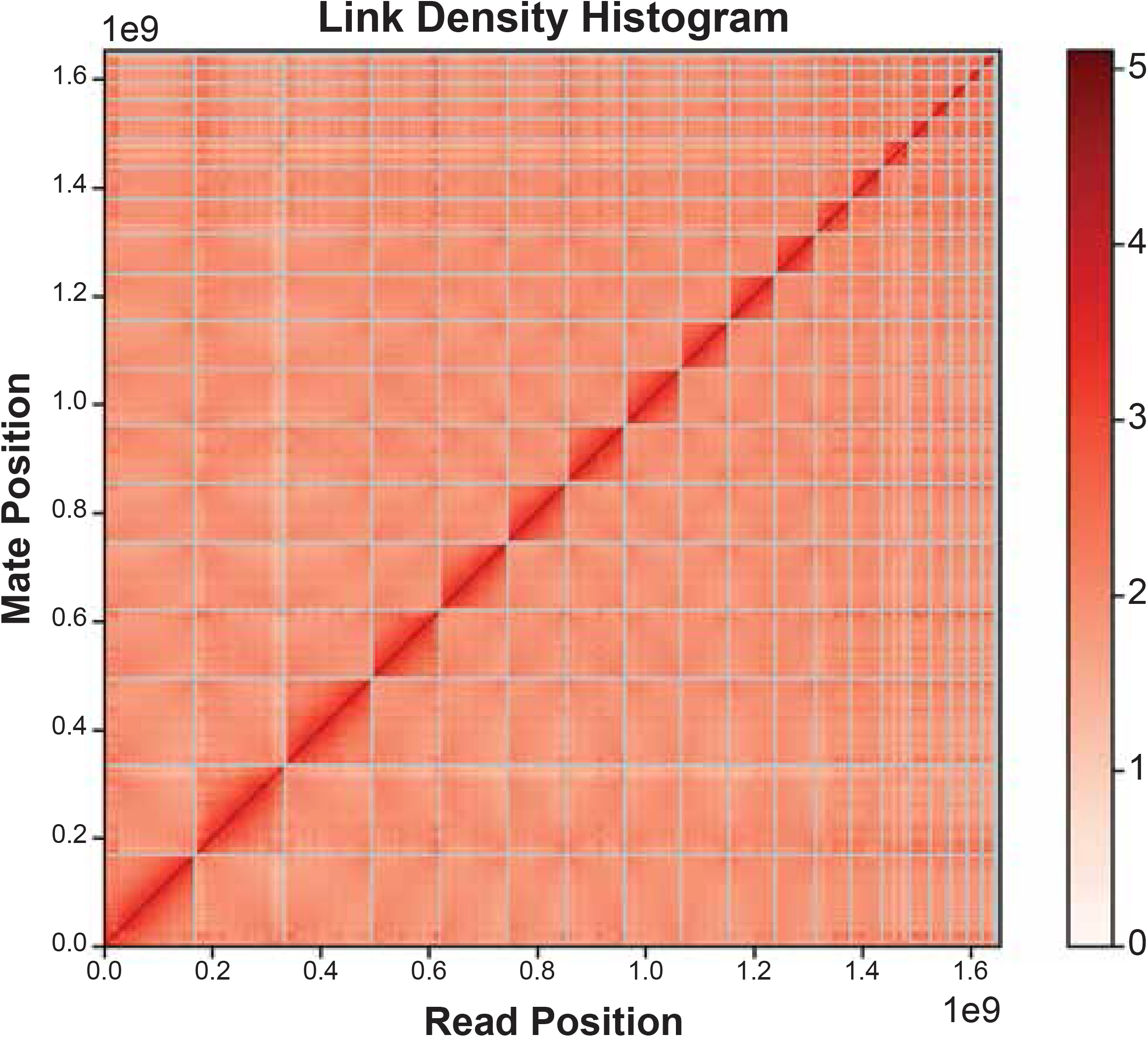
Hi-C contact plot of 19 chromosomal scaffolds, along with unassigned scaffolds.

The repetitive content consisted of 40.41% of the *C. ciliatus* genome, with a total length of 663.95 Mbp (Table 5). DNA transposons consisted of 1.39%, while LINE, SINE, and LTR transposons consisted of 14.75%, 6.42%, and 1.08% of the genome respectively. The de novo gene prediction resulted in a total of 30,780 protein coding genes (Table 6), and of the identified genes, 20,429 (66.37%) had an AED score ≤ 0.5 (Supplementary Material Figure 1).

**Table 5.** Summary of Repetitive Content and Transposable Elements of *C. ciliatus* genome assembly.

| Type | Number of Elements | Length (bp) | % of genome |
| --- | --- | --- | --- |
| DNA | 173526 | 23044151 | 1.39 |
| LINE | 789302 | 243834212 | 14.75 |
| SINE | 652803 | 106143112 | 6.42 |
| LTR | 58109 | 17890933 | 1.08 |
| Other | 889 | 381767 | 0.02 |
| Unknown | 1902327 | 272947662 | 16.51 |
| Small RNA | 2588 | 3815594 | 0.23 |
| Satellites | 1039 | 952457 | 0.06 |
| Simple repeats | 292097 | 14087986 | 0.85 |
| Low complexity | 25303 | 1434776 | 0.09 |
| Total | 3897983 | 684532650 | 40.41 |

**Table 6.** De Novo Gene Prediction Metrics.

|  |  |
| --- | --- |
| Total Number of Genes | 30,780 |
| Total coding region (bp) | 39,760,070 |
| Average length of genes (bp) | 1,291.75 |
| Number of single-exon genes | 1,461 |

## Data Validation and Quality Control

BUSCO v5.7.1 [24] was used to assess the quality and completeness of the *C. ciliatus* genome. BUSCO analysis was performed using the eukaryota_odb10 dataset. The *C. ciliatus* genome captured 99.6% of BUSCOs in the eukaryota_od10 dataset (Table 7), indicating the high completeness of the assembly.

**Table 7.** BUSCO analysis summary of *C. ciliatus* genome.

| BUSCO Category | Value | Percent (%) |
| --- | --- | --- |
| Complete BUSCOs | 254 | 99.6 |
| Complete and single-copy BUSCOs | 249 | 97.6 |
| Complete and duplicated BUSCOs | 5 | 2 |
| Fragmented BUSCOs | 1 | 0.4 |

### Availability of supporting data

Supporting datasets, including annotation are available at GigaDB. Raw sequencing reads and OmniC Library reads have been deposited in the SRA (Sequence Read Archive) database under Bioproject ID PRJNA1091669. RNA-Seq reads have been deposited under BioProject ID PRJNA1128839. This Whole Genome Shotgun project has been deposited at DDBJ/ENA/GenBank under the accession JBBPXQ000000000 and Biosample accession number is SAMN40604022. The version described in this paper is version JBBPXQ010000000.

## Authors’ contributions

TL conceived and supervised the project and provided the crested gecko samples. MG analyzed genome assembly performed repeat annotation and BUSCO analysis. MG drafted the manuscript. TL revised the manuscript. ZP maintained animal colonies. All authors read and approved the final manuscript. All authors read and approved the final manuscript.

## List of Abbreviations

BUSCO: Benchmarking Universal Single-Copy Orthologue
RNA-Seq: RNA Sequencing
BLASTN: Basic Local Alignment Search Tool (for nucleotides)
bp: base pairs
Mbp: Mega base pairs
TE: transposable element
hmwDNA: high molecular weight DNA
AED: Annotation Edit Distance

## Acknowledgements

We would like to acknowledge funding from NIH R01GM115444 and support from the CIRM COMPASS Award (EDUC5-13853). Special thanks to Dr. Andrew McMahon, the Department of Stem Cell Biology and Regenerative Medicine, and the Eli and Edythe Broad Center for Regenerative Medicine and Stem Cell Research at University of Southern California for genome sequencing support.

