## Supplementary Material Table 1; Supplementary Material Figure 1 for "Chromosome-level genome assembly and annotation of the crested gecko, *Correlophus ciliatus*, a lizard incapable of tail regeneration"

**This PDF file includes:**

Supplementary Material Table 1

Supplementary Material Figure 1

**Supplementary Material Table 1: Distributions, availabilities within American and European pet trades, and conservation statuses of the fourteen described geckos species lacking tail regenerative capabilities.**

| **Species** | **Distribution** | **Availability** | **Conservation Status (IUCN 3.1)** |
| --- | --- | --- | --- |
| *Correlophus ciliatus* | New Caledonia | ++++++ | Vulnerable |
| *Correlophus belepensis* | New Caledonia |  | Critically Endangered |
| *Nephrurus amyae* | Australia | +++ | Least Concern |
| *Nephrurus asper* | Australia | +++ | Least Concern |
| *Nephrurus sheai* | Australia | ++ | Least Concern |
| *Uroplatus ebenaui* | Madagascar | +++ | Vulnerable |
| *Uroplatus fetsy* | Madagascar |  |  |
| *Uroplatus fiera* | Madagascar | + |  |
| *Uroplatus finaritra* | Madagascar | + |  |
| *Uroplatus fotsivava* | Madagascar |  |  |
| *Uroplatus kelirambo* | Madagascar |  |  |
| *Uroplatus malama* | Madagascar |  | Vulnerable |
| *Uroplatus phantasticus* | Madagascar | +++ | Least Concern |

**Supplementary Material Figure 1:** The frequency of annotation edit distance (scores) for the *Correlophus ciliatus* assembly. Annotation edit distance (AED) is a general measure of how well the predicted gene is supported by external evidence (UniProt protein and mRNA sequences). AED score ranges from 0 to 1 and a

lower score represents more evidence support for the gene. AED is calculated for every gene. The

AED cumulative frequency graph above provides an overview of the quality of the gene annotation.

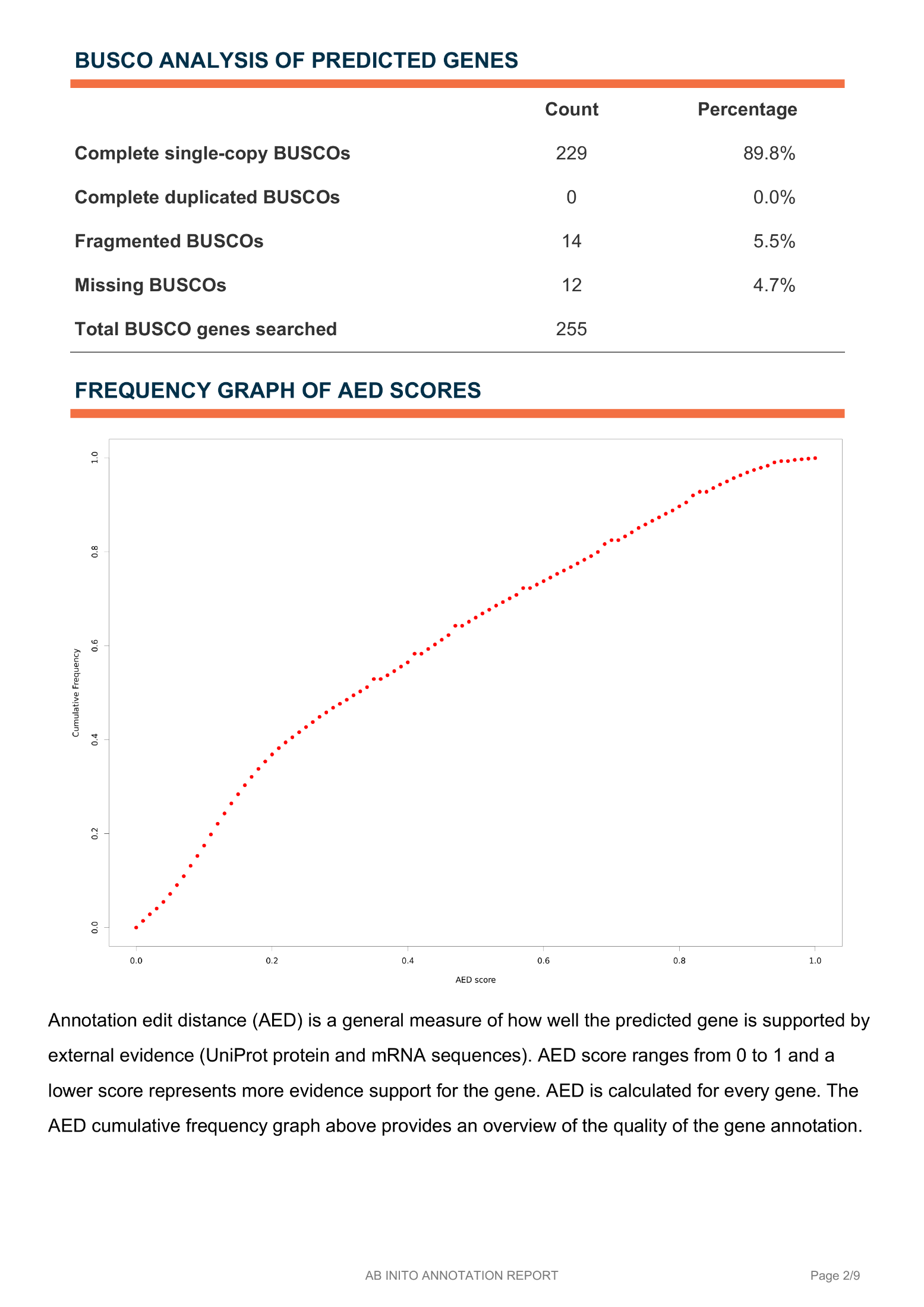
